# Protein Kinase D promotes activity-dependent AMPA receptor endocytosis in hippocampal neurons

**DOI:** 10.1101/2020.04.09.033514

**Authors:** Carlos O. Oueslati Morales, Attila Ignácz, Norbert Bencsik, Anikó Erika Rátkai, Wolfgang S. Lieb, Stephan A. Eisler, Attila Szűcs, Katalin Schlett, Angelika Hausser

**Affiliations:** Membrane Trafficking and Signalling Group, Institute of Cell Biology and Immunology, University of Stuttgart, Stuttgart, Germany; Neuronal Cell Biology Group, Department of Physiology and Neurobiology, Eötvös Loránd University, Budapest, Hungary; Stuttgart Research Center Systems Biology, University of Stuttgart, Stuttgart, Germany

**Keywords:** AMPA receptor, GluA1, endocytosis, chemically induced LTD, PKD

## Abstract

AMPA type glutamate receptors (AMPARs) mediate the majority of fast excitatory neurotransmission in the brain. The continuous trafficking of AMPARs into and out of synapses is a core feature of synaptic plasticity, which is considered as the cellular basis of learning and memory. The molecular mechanisms underlying the postsynaptic AMPAR trafficking, however, are still not fully understood. In this work, we demonstrate that the Protein Kinase D (PKD) family promotes basal and activity-induced AMPAR endocytosis in primary hippocampal neurons. Pharmacological inhibition of PKD increased synaptic levels of GluA1-containing AMPARs, slowed down their endocytic trafficking and increased neuronal network activity. By contrast, ectopic expression of constitutive active PKD decreased the synaptic level of AMPARs, while increasing their co-localization with early endosomes. Our results thus establish an important role for PKD in the regulation of postsynaptic AMPAR trafficking during synaptic plasticity.

## Introduction

Synaptic plasticity describes a process where synaptic strength is changed in an activity-dependent manner within the brain. The most widely investigated forms of long-term synaptic changes include long-term potentiation (LTP) and long-term depression (LTD), which are thought to present the cellular mechanisms of learning and memory (Malinow & Malenka 2002; Huganir & Nicoll 2013). One of the key elements in the control of synaptic strength is the dynamic alteration in the number and composition of the ionotropic AMPA-type glutamate receptors (AMPARs) within the postsynaptic membrane (Malinow & Malenka 2002; Huganir & Nicoll 2013; Chater & Goda 2014).

AMPARs are tetramers composed of four different subunits, GluA1-GluA4. Binding of glutamate allows the opening of the ion channel and the subsequent influx of Na^+^ (and potentially Ca^2+^) and efflux of K^+^, causing membrane depolarization. Thus, the main function of AMPARs is to mediate excitatory neurotransmission in the brain (Chater & Goda 2014; Diering & Huganir 2018). The heterotetramer GluA1/2 is the most abundant variant followed by GluA2/3, while the GluA4 subunit is expressed mainly during early development and is present only at low levels in the adult brain (Henley & Wilkinson 2013; Lu *et al*. 2009; Zhu *et al*. 2000). GluA1-containing AMPARs leave the endoplasmic reticulum rapidly and are trafficked towards the synaptic plasma membrane. Plasma membrane insertion is thought to happen mostly within the extra- or perisynaptic regions, followed by lateral movement of AMPARs into the synapse. Within the synaptic membrane, AMPARs are stabilized by postsynaptic density (PSD) scaffolding proteins in an activity-dependent manner (Greger *et al*. 2002; Henley & Wilkinson 2013; Makino & Malinow 2009; Pick & Ziff 2018). Synaptic AMPARs are constitutively recycled through endocytic trafficking pathways: AMPARs are internalized from the cell surface at perisynaptic endocytic zones. Subsequently, AMPARs are trafficked to early endosomes and either recycled back to the plasma membrane or transported to dendritic lysosomes and degraded (Blanpied *et al*. 2002; van der Sluijs & Hoogenraad 2011; Luscher *et al*. 1999; Ehlers 2000). During LTD and LTP, endocytotic and exocytotic AMPAR trafficking is imbalanced, resulting in reduced and enhanced numbers of AMPARs at the synapse, respectively (Carroll *et al*. 1999; Ehlers 2000; Hayashi *et al*. 2000; Shi *et al*. 1999).

The protein kinase D (PKD) family of serine/threonine kinases belongs to the calcium/calmodulin-dependent protein kinases (CaMKs) superfamily, and comprises three isoforms in mammals: PKD1, PKD2 and PKD3, all of which are expressed in neurons from a very early embryonic stage (Ellwanger & Hausser 2013; Oster *et al*. 2006). PKDs are recruited to the plasma membrane or to different organelles, such as the Golgi complex, by binding to diacylglycerol. Upon activation, they control different intracellular processes such as vesicle fission from the Golgi complex and rearrangement of the actin cytoskeleton (reviewed in (Olayioye *et al*. 2013; Reinhardt *et al*. 2020)). In breast cancer cells, PKD-mediated phosphorylation of Rabaptin-5, a binding partner for both Rab4 and Rab5, is required to shunt integrin αvβ3 to the short recycling loop, indicating a role for the kinase in endocytic trafficking processes (Christoforides *et al*. 2012; Woods *et al*. 2004). In neurons, PKD plays a key role in the establishment and maintenance of polarity through controlling the integrity of the Golgi complex and trans-Golgi network-derived sorting of vesicles (Bisbal *et al*. 2008; Czondor *et al*. 2009; Yin *et al*.2008). Furthermore, we recently demonstrated a role for PKD in learning and memory through stabilization of plasticity-induced actin rearrangements (Bencsik *et al*. 2015). However, so far PKD has not been linked to AMPAR trafficking in neuronal cells.

Here, we identify PKD as a regulator of basal and activity-mediated GluA1 endocytosis in cultured hippocampal neurons. We provide evidence that PKD is activated downstream of chemically induced GluA1 endocytosis. Inhibition of PKD increases surface and synaptic levels of GluA1 and consequently, neuronal network activity. We further show that expression of constitutive active PKD (caPKD) decreases the synaptic amount of GluA1 while increasing the localization of GluA1 at early endosomes. Importantly, PKD inhibition blocks the endocytosis of GluA1 upon chemically evoked LTD (cLTD). Combined, our data identify a previously unknown role of the PKD family in the regulation of AMPAR endocytosis in cultured hippocampal neurons.

## Material and methods

### Animal handling

Female pregnant CD1 mice (Charles River) were kept alone in type 2 cages (Zoonlab #3010010) in the animal facility of the Institute of Cell Biology and Immunology, University of Stuttgart; or of the Biological Institute, Eötvös Loránd University, at 22 ± 1°C with 12 h light and dark cycles and ad libitum access to water and food. The animals were maintained and handled in accordance with the Guidelines for Accommodation and Care of Animals, according to the European Convention for the Protection of Vertebrate Animals Used for Experimental and Other Scientific Purposes.

### Cell culture

Primary cultures of embryonic hippocampal cells were prepared from CD1 mice on embryonic day E17.5-E18.5. In brief, E 17.5-18.5 pregnant female CD1 mice were sacrificed by means of CO_2_ intoxication, and subsequently the hippocampi were aseptically removed from the skull of the embryos. Tissue was then freed from meninges and incubated in 0.05% trypsin-EDTA (ThermoFisher Scientific #15400-054) solution with 0.05% DNAseI (Sigma-Aldrich #D5025) in PBS for 15 min at 37°C. After a brief centrifugation, cells were disaggregated in NeuroBasal culture medium (ThermoFisher Scientific, #21103049) supplemented with B27 (ThermoFisher Scientific, #17504044), 5% fetal bovine serum (PAN Biotech #P30-3309), 0.5 mM GlutaMAX (ThermoFisher Scientific, #35050-038), 40 μg/ml gentamycin (Sigma, #G1397) and 2.5 μg/ml amphotericin B (Gibco, #15290-026) and filtered through a sterile polyester mesh with 42-μm pore size (EmTek, Budapest, Hungary). Cell number was determined by trypan blue exclusion, and cells were seeded in supplemented NeuroBasal on poly-L-lysine-coated (PLL, Sigma #5899), 6 well plates (Greiner Bio-One, #657160) or 35 mm petri dishes (Greiner Bio-One, #627160) at a density of 4 x 10^5^ cells/well. For imaging, cells were seeded on PLL/laminin-coated (Sigma; #L2020) glass coverslips (Carl Roth, #YX03.1) in 24-well plates or on glass bottom petri-dishes (Greiner Bio-One, #627870) at a density of 1.15 x 10^5^ cells/well. Five days after plating, half of the original NeuroBasal culture medium (ThermoFisher Scientific, #21103049) supplemented with B27 (ThermoFisher Scientific, #17504044) was changed to Brainphys (Stemcell technologies, #05790) supplemented with SM1 (Stemcell technologies, #05711), gentamycin and amphotericin B, containing 10 μM cytosin-arabinofuranoside (Sigma, #C6645). The cells were cultivated for 13 days at 37°C in 5% CO_2_, and one half of the Neurobasal media was exchanged by fresh supplemented Brainphys every 3 days. Chemically-induced Long Term Depression (cLTD) was induced by treating the neuronal cultures with 50 μM NMDA (Sigma, #M3262) for 5 min in conditioned medium. Then, cells were changed to a NMDA-free medium, and incubated for 15 min before being further processed. Agonist-induced AMPAR endocytosis was performed by treating the neuronal cultures with 100 μM S-AMPA (Hello Bio, #HB0052) and 50 μM D-AP5 (Tocris #3693) for 2 minutes in conditioned medium. Afterwards, cells were changed to an AMPA- and D-AP5-free medium and incubated for 10 minutes before being further processed.

### Chemical treatments, cell surface staining and biotinylation

PKD activity was blocked with CRT0066101 (CRT, Tocris #4975) (Harikumar *et al*. 2010) diluted in dimethyl sulfoxide (DMSO), at a final concentration of 2 μM. For surface biotinylation of endogenous GluA1, 13-day-old neuronal cultures were placed on ice (13 days in vitro; DIV13), washed twice with calcium- and magnesium-containing phosphate-buffered saline (PBSCM, ThermoFisher Scientific, #14040-091) and incubated with 1.5 mg/ml sulfo-NHS-SS-biotin (ThermoFisher Scientific, #21331) diluted in PBSCM for 15 min. Afterwards, cultures were washed twice with 20 mM glycine (Roth, #HN07.1) in PBSCM for 7 min, to bind free biotin; before being finally lysed. To visualize cell surface GluA1, we modified an assay from a published protocol (Sziber *et al*. 2017). In brief, neuronal cultures were incubated with an N-terminal-specific GluA1 antibody (1:100, Millipore, #MAB2263, RRID:AB_11212678) in conditioned media for 10 min at 37 °C and 5% CO_2_ in a humidified atmosphere. Afterwards, cells were washed with PBS and fixed with 4% paraformaldehyde (PFA) in PBS for 10 min at room temperature (RT) without permeabilization. After 1 h of blocking with 5% FBS and 0.1% NaN3 in PBS, a secondary anti-mouse antibody labelled with Alexa 546 (1:500, Thermo Fisher Scientific, #A11030, RRID:AB_2534089) was applied to the cells for 1h at RT. Cells were washed and fixed again with 4% PFA in PBS for 10 min and further processed for Shank2 staining (see section Immunostaining and confocal laser scanning microscopy). Samples were mounted in ProLong Gold Antifade Mountant (ThermoFisher Scientific, #P36930) and analyzed by confocal laser scanning microscopy.

To analyze the amount of endocytosed AMPAR, biotinylation assays of receptor internalization were employed as described (Ehlers 2000). In brief, protein lysosomal degradation was inhibited with 100 μg/ml Leupeptin (Tocris, #1167) for 1h; then cells were placed on ice and incubated with 1.5 mg/ml Sulfo-NHS-SS-biotin (ThermoFisher Scientific, #21331) for 1 h. Subsequently, cells were transferred to biotin-free medium and treated with 100 μM S-AMPA and 50 μM D-AP5 for 2 min. Then, cells were returned to an AMPA- and D-AP5 free medium for 10 min to allow for endocytosis, subsequently placed again on ice and washed twice with cold PBSCM. Remaining surface biotin was cleaved by incubation with glutathione cleavage buffer (50 nM L-Glutathione (Sigma #G6013) 75 mM NaCl, 10 mM EDTA, 1% BSA, pH 8.6) for 15 min. Finally, cells were washed twice and further processed for protein extraction.

### Protein extraction, biotin pulldowns and Western Blotting

Neuronal cells were harvested in cold lysis buffer (1% Triton X-100, 20 mM Tris pH 7.5, 150 mM NaCl, 1mM EDTA, 1 mM EGTA; 4°C) supplemented with protease (Roche, #11697498001) and phosphatase inhibitors (Roche, #04906845001), and subsequently centrifugated at 13000 x g for 10 min. For the biotin pulldowns, equal amounts of cell lysate were incubated with avidin-coated agarose beads (Thermo Scientific, #29201) for 90 min at 4°C and washed 4 times with lysis buffer. Equal amounts of protein were run on NuPage Novex 4-12% Bis-Tris gels (Thermo Fisher Scientific, #NP0322BOX) and blotted onto nitrocellulose membranes using the iBlot device (Thermofisher scientific, #IB1001). Membranes were blocked for 30 min with 0.5% (v/v) blocking reagent (Roche, #11096176001) in PBS containing 0.05% Tween 20 and 0.1% NaN3, and subsequently incubated with primary antibodies overnight at 4°C (anti-PKD1 (1:1000, Cell Signaling Technology #2052, RRID:AB_2172539), anti-pS916PKD (1:1000, CST #2051, RRID:AB_330841), anti-actin (1:1000, Sigma-Aldrich #A4700, RRID:AB_476730), anti-GluA1 (1:1000, Millipore #MAB2263, RRID:AB_11212678), anti-pS831GluA1 (1:1000, Cell Signalling Technology #75574, RRID:AB_2799873), anti-GFP (1:2000, Roche #11814460001, RRID:AB_390913)). The next day, membranes were washed and incubated with HRP-conjugated secondary anti-mouse and anti-rabbit antibodies at RT for 1 h (1:10000, Jackson ImmunoResearch Labs #115-035-062, RRID:AB_2338504 or #111-035-144, RRID:AB_2307391) before being washed again and finally visualized using the SuperSignal West Pico PLUS or the SuperSignal West Dura Extended Duration substrates (Thermo Fisher Scientific, #34580 or #34076). Proteins were visualized using the Amersham Imager 600, an enhanced chemiluminescence detection system (Thermo Fisher Scientific, Waltham, MA, USA). Quantitative Western Blotting chemiluminescence was detected at a depth of 16-bit in the linear detection range of the device, equipped with a 3.2-megapixel super-honeycomb CCD camera fitted with a large aperture f/0.85 FUJINON lens. Special care was taken not to overexpose in order to guarantee accurate quantifications. Densitometry was performed using Image Studio Lite 5.2 (Image Studio Lite, RRID:SCR_013715). For each protein, the integrated density of the signal was measured, corrected for background signals and adjusted to loading controls.

### Transfection and expression constructs

On DIV12, neuronal cultures were transfected using Lipofectamine 2000 (Invitrogen, #11668-019) according to the manufacturer’s instructions. In brief, 0.5 μg of plasmid DNA was mixed with Lipofectamine 2000 in a 1 μg: 2 μl ratio. Medium was changed 4 h after transfection to the original cultivation medium and cells were incubated overnight. The enhanced green fluorescent protein (EGFP) vector was obtained from Clontech Europe, whereas the pcDNA3-mRuby2 was a gift from Michael Lin (Addgene plasmid #40260; http://n2t.net/addgene:40260; RRID:Addgene_40260) and pCI-SEP-GluR1 was a gift from Robert Malinow (Addgene plasmid #24000; http://n2t.net/addgene:24000; RRID:Addgene_24000) (Kopec *et al*. 2006). caPKD1-EGFP and caPKD1-Flag have already been described elsewhere (Hausser *et al*. 2002).

### Immunostaining and confocal laser scanning microscopy

Hippocampal cultures were fixed on DIV13 for 20 min with 4% PFA in PBS. After washing with PBS, cells were permeabilized with 0.1% Triton X-100 in PBS for 5 min. Cells were again washed with PBS and blocked with 5% FBS and 0.1% NaN3 in PBS for 1 h at RT. Subsequently, the corresponding primary antibody (anti-Shank2 (1:2000, Synaptic Systems #162 204, RRID:AB_2619861), anti-GluA1 (1:500, Millipore #MAB2263, RRID:AB_11212678), anti-EEA1 (1:100, Cell Signalling Technology #3288, RRID:AB_2096811)) was diluted in blocking buffer and incubated for 1.5 h at RT. Cells were washed with PBS, and Alexa-Fluor-(488, 546 or 633) labelled secondary antibodies (ThermoFisher Scientific #A-11029, RRID:AB_2534088; #A-11030, RRID:AB_2534089 and #A-21052, RRID:AB_2535719, respectively) were diluted 1:500 in blocking buffer and incubated for 1 h at RT. Finally, samples were mounted in ProLong Gold Antifade. All samples were imaged using a confocal laser scanning microscope (LSM 710, Carl Zeiss) equipped with either a Plan Apochromat 63x/1.40 DIC M27 or an alpha Plan-Apochromat 100x/1.46 Oil DIC objective (Carl Zeiss), using sequential excitation with an 488 nm Argon laser, an 561 DPSS laser or an 633 nm HeNe laser. Image acquisition for the quantitative measurement of GluA1 intensity was done as follows: z-stacks of 0.32 μm intervals were acquired throughout the selected neuronal branches of at least 20 μm in length. Image processing and analysis were performed with either ZEN blue (ZEN Digital Imaging for Light Microscopy, RRID:SCR_013672) or with ImageJ (Image J, RRID:SCR_003070). Regions of interests were selected manually according to clear Shank2 immunopositivity at the plasma membrane along the shaft or within dendritic spines. Mean pixel intensity values of the GluA1 channel within the selected regions of interests were measured, background corrected and normalized to the staining intensity in the control condition. In order to analyze the co-localization between GluA1 and EEA1 signals, single images of selected neuronal branches were acquired. In all experiments, laser power was set so that there would be no saturation and maintained constant throughout the analyses of different samples from the same experiment. Image processing and analysis was performed with ZEN blue and ImageJ. The Mander’s overlap coefficient of GluA1 with EEA1 was calculated and used as a measure of co-localization using the ImageJ plugin JACoP.

### Fluorescence recovery after photobleaching (FRAP)

To investigate AMPAR trafficking to and from the plasma membrane, neuronal cultures were transfected with the super ecliptic pHluorin-tagged GluA1 (GluA1-SEP) and mRuby2 plasmids in a 5:3 ratio and incubated overnight. On the next day, culture medium was changed to pre-heated live cell imaging buffer containing 142 mM NaCl, 5.4 Mm KCl, 1.8 mM CaCl_2_, 1 mM NaH_2_PO_4_, 0.8 mM MgSO_4_, 5 mM glucose and 25 mM HEPES with a pH of 7.4. The diffusion mobility of the GluA1-containing AMPARs was analyzed at 37°C and 5% CO_2_ on a Zeiss Axio Observer Spinning Disc Microscope, using an alpha Plan-Apochromat 100x/1.46 Oil DIC objective and a Photometrix Evolve 512 EMCCD camera for image acquisition. Photobleaching was executed with a UGA-42 firefly photomanipulation system equipped with a 100 mW 473 nm laser (Rapp OptoElectronic, Germany). Cells were initially treated with 2 μM of CRT for 10 min, then selected dendritic spines were bleached with a high-intensity laser light (473 nm line, 10% laser power). Fluorescence intensity in the bleached areas was measured over time both before and after the bleaching event, taking images every 30 seconds for approx. 25 min. Intensity values were corrected with the background intensity values and normalized to the unbleached region. Fitting of the curves was performed with a one-phase exponential equation *Y* = *Y*0 +(*Plateau* − *Y*0) * (1 − *exp*(−*K* * *x*)); Prism 8, (GraphPad Prism, RRID:SCR_002798).

### MEA recording and analysis

For extracellular voltage recording, hippocampal cell cultures were seeded into PLL/laminin coated Axion 24-well BioCircuit (M384-tMEA24W) and 48-well CytoView (M768-tMEA-48B) MEA plates in a density of 1.5×10^5^ and 4×10^4^ cells per well, respectively. Five days after plating, half of the culture medium was changed to Brainphys supplemented with SM1, gentamycin and amphotericin B, and treated with cytosin-arabinofuranoside. At 12-14 days in vitro, baseline neuronal activity was recorded for 5 minutes by the Maestro Pro recording system (Axion Biosystems) or by a custom-made recording hardware (APPERCELL Biotech Ltd., Hungary), followed by DMSO, CRT and cLTD treatments as described above. 5 minutes long recordings were analysed between 10 to 15 minutes and 90 to 95 minutes following the corresponding treatments. Action potentials from raw voltage recordings were detected by the Axion AxIS Navigator software or by the custom-written software “Neuroexpress” developed by A. Szűcs^1^. Spike arrival time data were analysed by the NeuroExpress software using a variety of tools including inter-spike time intervals (ISI) and burst cycle periods (BCP) as measures of the general intensity of neuronal activity. Bursts were defined as clusters containing at least 3 action potentials with adjacent spikes occurring within 460 ms. Burstiness, a dimensionless parameter was used to describe the overall clustering and separation of burst episodes in the firing pattern. To calculate this parameter, first we identified the smaller of the 2 interburst intervals either preceding or following the actual burst event. Next, we determined the arithmetic mean of the smallest and longest intraburst ISI for the burst. Burstiness was then calculated by dividing the smaller interburst interval by the mean ISI and subtracted 1 from the resulting fraction. Burstiness, therefore, was obtained for each burst event in the spike train. This parameter yields zero for periodic firing (pacemaker pattern), while increasingly higher values are obtained when the firing becomes more clustered.

### Statistical analysis of the data

For statistical evaluation, the normality of the data was routinely tested using the Shapiro-Wilk normality test. For the comparison of two groups, a t test was employed, whereas to compare three or more groups, a one-way ANOVA was performed followed by a post-hoc test. No test for outliers was conducted and no data points were excluded. Statistical analysis was tested employing GraphPad Prism 8 software. Data are displayed as mean ± SEM, mean ± 95% confidence interval (CI) or Tukey’s box-and-whiskers plots.

### Ethical approval

All applicable international, national, and/or institutional guidelines for the care and use of animals were followed. The animal experiments conducted in Stuttgart and Budapest were reviewed and approved by the Regierungspräsidium Stuttgart and by the Animal Ethics Committee of Eötvös Loránd University (approval number: PEI/001/1108-4/2013 and PEI/001/1109-4/2013), respectively. This article does not contain any studies with human participants performed by any of the authors.

## Results

### AMPAR basal trafficking is regulated by PKD

AMPAR surface expression and recycling are highly dynamic, with a half-time in the range of minutes (Henley & Wilkinson 2013). To address the role of PKD in basal AMPAR trafficking, we thus subjected hippocampal neurons to an acute, short-term PKD inhibition (10 min) using CRT0066101 (referred to as CRT), a potent and selective pan-inhibitor of all PKD isoforms (Harikumar *et al*. 2010; Borges *et al*. 2015; Lieb *et al*. 2020; Zhang *et al*. 2017). In order to prove that this short-term treatment with CRT effectively inhibits PKD activity, we analyzed autophosphorylation of PKD at serine 916 (pS916PKD) immediately following CRT treatment, as an indicator of kinase activity (Matthews *et al*.1999) along with the total levels of PKD (Figure 1a). In line with our previous results (Czondor *et al*.2009), we show that endogenous PKD is active in cultured hippocampal neurons under basal conditions. We further detect a strong and significant decrease in the relative level of pS916PKD in CRT treated cells compared to the DMSO treated control (Figure 1a, b), confirming the efficacy of the inhibitor under these conditions.

**Fig. 1.**
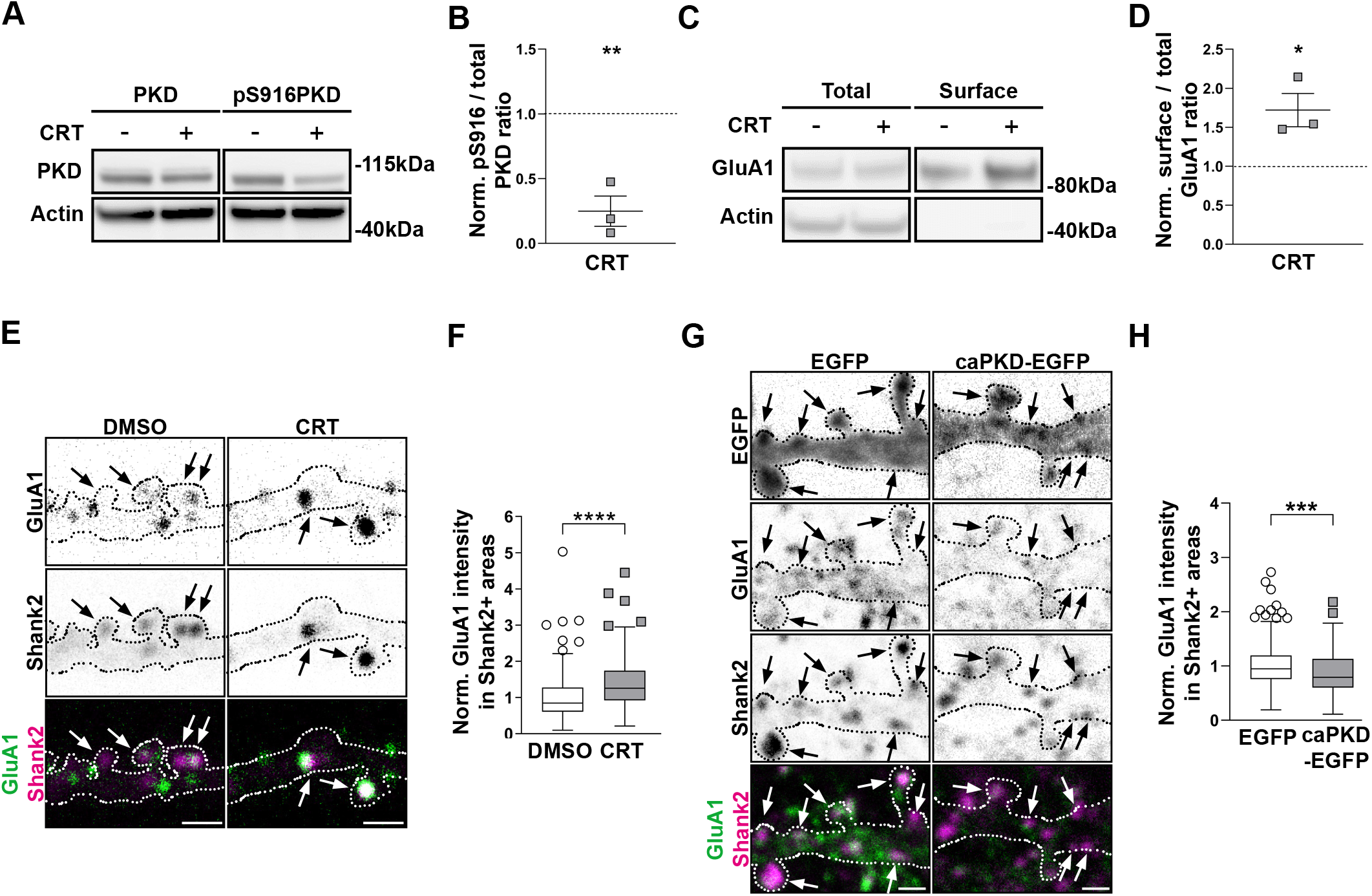
PKD regulates basal AMPAR trafficking. Representative Western Blots obtained from hippocampal neuronal cultures treated with DMSO (-) or CRT displaying PKD and pS916PKD levels (a) and total and surface GluA1 levels upon biotinylation (c). Actin served as a loading control and is absent from the precipitated samples. (b and d) Quantification of data shown in (a) and (c), respectively, using densitometry analysis. Data from CRT-treated cultures are presented as mean line density ± SEM normalized to the control cultures; each dot represents one independent culture. Statistical comparison was performed by unpaired two-tailed t-test. The dotted line indicates the control level. (e and g) Representative inverted single-channel and colored merged pictures of neuronal dendritic branches treated with DMSO or CRT (e) or expressing EGFP or caPKD-EGFP (g) stained for the extracellular N-terminal domain of GluA1 and the PSD marker Shank2. Arrows point to Shank2-positive areas included in the analysis. The EGFP signal is not included in the colored merged image. (f and h) Quantification of pictures shown in (e) and (g), respectively. Data were normalized for the DMSO (f) or EGFP control (h). The boxplots show the results of three independent cultures with 11 (DMSO), 12 (CRT), 17 (EGFP) and 12 (caPKD) neurons analyzed. Center lines show the medians; box limits indicate the 25th and 75th percentiles; whiskers extend 1.5 times the interquartile range from the 25th and 75th percentiles, outliers are represented by dots. The number of investigated sample points (Shank2+ areas) is n=171 and 273 (f) or n=220 and 223 (h), respectively. The significance of differences was analyzed by a Mann-Whitney test (two-tailed).

Next, to assess the surface expression of GluA1-containing AMPARs in hippocampal neurons treated with CRT or the solvent DMSO, we utilized cell surface protein tagging using sulfo-NHS-SS-biotin. After cell lysis, biotinylated proteins were precipitated and the total expression and surface levels of GluA1 were detected by Western Blot analysis (Figure 1c). Our data clearly indicate that already a short-term PKD inhibition significantly increased the relative amount of surface GluA1 compared to the DMSO treatment (Figure 1d).

Under basal conditions, AMPARs have a high turnover within spines (Passafaro *et al*. 2001; Shi *et al*.2001) and are internalized in the vicinity of synapses (Rosendale *et al*. 2017). To investigate whether short-term PKD inhibition affects the amount of GluA1 within the close vicinity of the postsynaptic membrane, we treated hippocampal cultures with CRT or DMSO, and subsequently incubated the cells with an antibody detecting the extracellular N-terminal domain of GluA1. Cells were then fixed and incubated with a fluorescently labelled secondary antibody without permeabilization to visualize GluA1 subunits within the plasma membrane. Subsequently, cell membrane was permeabilized and cells were additionally immunostained for Shank2, a marker of the PSD (Naisbitt *et al*. 1999) (Figure 1e). Notably, our analysis reveals that inhibition of PKD significantly increased GluA1 intensity within Shank2-positive areas compared to the control (Figure 1f). Vice versa, transfection of hippocampal cells with a plasmid encoding a constitutively active S738/742E mutant form of PKD (caPKD-EGFP) significantly decreased the amount of GluA1 in Shank2 positive areas compared to cells transfected with EGFP as control (Figure 1 g, h).

To measure the impact of PKD inhibition on spontaneous neuronal activity, we cultivated hippocampal neurons on transparent Multi-electrode Array (MEA) plates containing 16 electrodes per well (see Figure 2a). Neurons at DIV14-15 were treated with DMSO or CRT for 10 min and subsequent extracellular voltage recordings were compared to baseline activity (Figure 2b). Short-term inhibition of PKD by CRT resulted in a significant decrease in interspike intervals compared to pre-treatment, basal values while spontaneous firing activity was not changed in DMSO-treated cultures (Figure 2c). Accordingly, when the effects of DMSO and CRT treatments were normalized to their corresponding baseline activity, CRT treatment significantly increased firing activity as interspike intervals were reduced (Figure 2d). In addition, 10 min CRT treatment significantly increased burstiness – a dimensionless parameter to describe the overall clustering of burst episodes - compared to basal values (Figure 2e) or to DMSO treatment (Figure 2f).

**Fig. 2.**
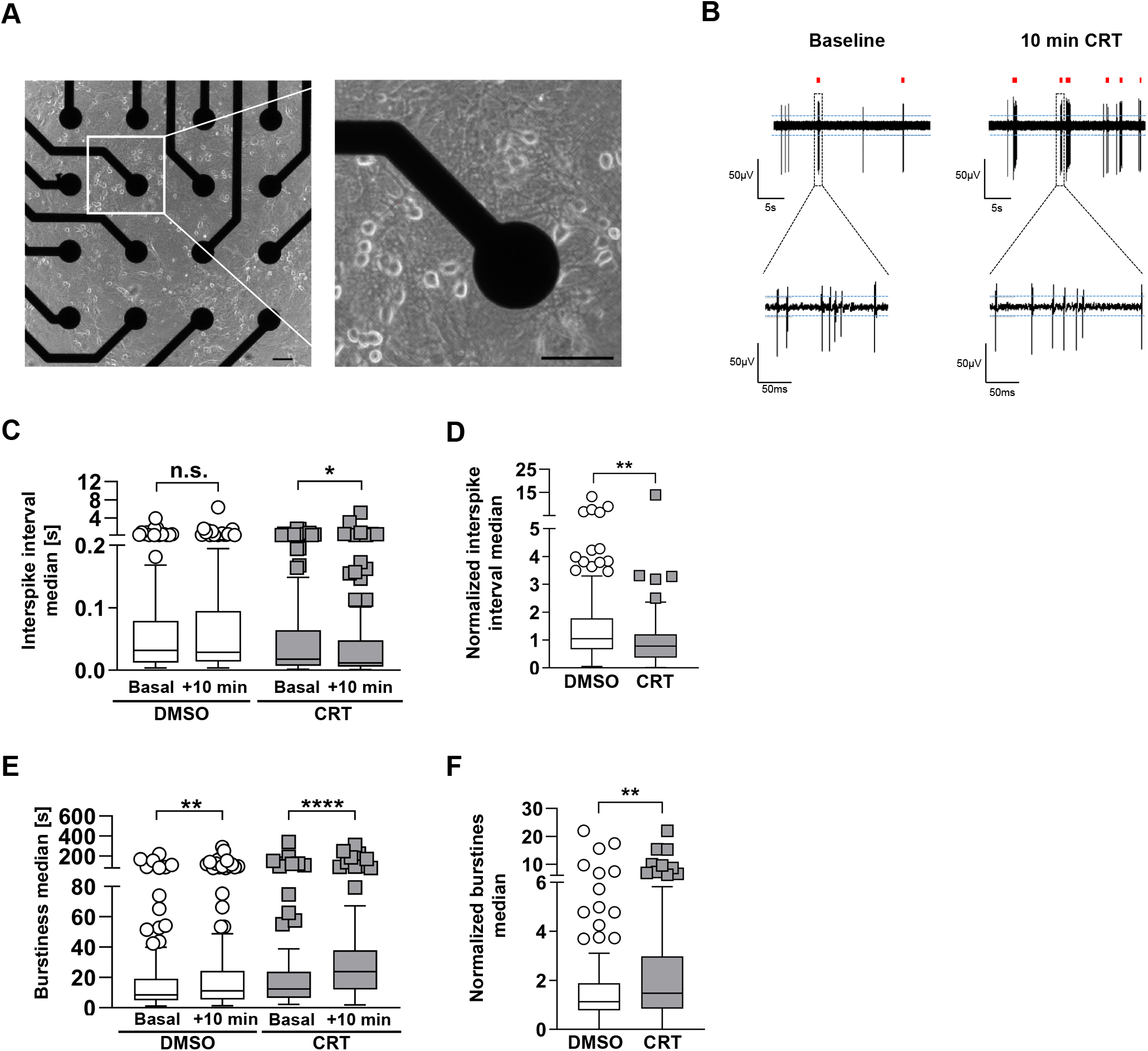
PKD regulates spontaneous neuronal network activity. (a) Representative phase contrast images of a MEA plate well at 13 days in vitro (DIV13) age. Scale bar 1 μm (left), 100 μm (right). (b) Short segments of raw voltage traces before and after CRT treatment. (c, e) Raw MEA data: Quantification of the raw interspike interval median (c) and the raw burstiness median (e) of neuronal cultures both before and after treatment with DMSO or CRT for 10 min. (d and f) Quantification of data shown in (b). The interspike interval median (d) and the burstiness median (f) were normalized for the DMSO controls. The boxplots show the results of three independent cultures. Center lines show the medians; box limits indicate the 25th and 75th percentiles; whiskers extend 1.5 times the interquartile range from the 25 th and 75th percentiles, outliers are represented by dots. The significance of differences was analyzed by a Mann-Whitney test (two-tailed) (d, f) or a Wilcoxon matched-pairs signed rank test (two-tailed) (c, e). *p<0.05, **p<0.01, ***p<0.001, ****p<0.0001.

Hence, our data provide compelling evidence that PKD activity regulates the amount and PSD localization of GluA1-containing AMPARs in synapses and decreases spontaneous network activity under basal conditions.

### PKD controls basal AMPAR endocytosis

To further investigate how PKD controls AMPAR trafficking, we measured fluorescence recovery after photobleaching (FRAP) of Super Ecliptic pHluorin (SEP)-tagged GluA1. SEP emits fluorescence under neutral pH conditions, allowing for the study of specific subpopulations of the tagged protein at the cell surface in relation to those fluorochromes which are internalized within vesicles (Kopec *et al*.2006). Cultures were co-transfected with mRuby2 to visualize transfected neuronal cells. Neurons were treated with either CRT or DMSO for 10 min, then the fluorescent signal within the dendritic spines of imaged dendritic branches was bleached and the recovery of the fluorescence intensity was measured for another 10 min (Figure 3a, b). To calculate the recovery half-time and the mobile fraction of GluA1-SEP, we additionally fitted the recovery curves with a one-phase exponential equation (Figure 3c). In DMSO treated neurons, photobleaching of spine SEP fluorescence to background levels was followed by a fast recovery (t½= 14.52 seconds with 95% CI of + 15.08 CI, Figure 3d) to 61.16% with a 95% CI of ± 3.4 of the original GluA1-SEP intensity after 183 seconds (Figure 3e). Upon PKD inhibition, GluA1-SEP signal within the spines recovered significantly slower (t½= 114.47 seconds with 95% CI of 77.67 and 38.26, Figure 3d), albeit to the same level as the control (Fig. 3b, c). Consequently, the mobile fraction of GluA1-SEP did not significantly change between the CRT and DMSO treated cultures (Figure 3e). The timescale of GluA1-SEP signal recovery depends on the net effects of lateral diffusion of plasma-resident, non-bleached GluA1-SEP subunits and on the rate of endocytic removal of bleached GluA1-SEP from the cell surface. Thus, our results suggest that PKD inhibition slows down the trafficking of cell surface GluA1-SEP.

**Fig. 3.**
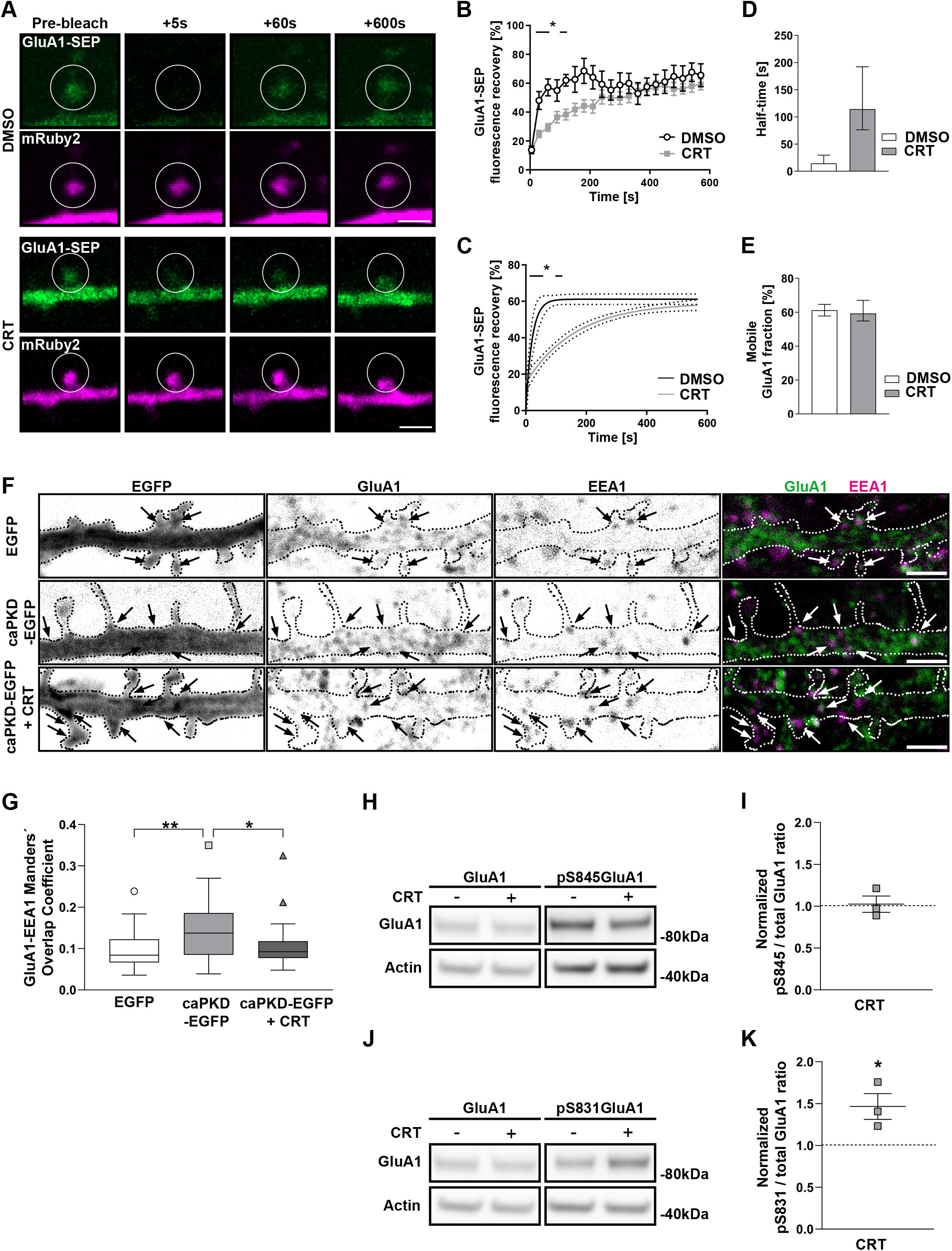
PKD specifically controls AMPAR endocytosis. (a) Representative pictures of neuronal dendritic spines expressing Super Ecliptic pHluorin-tagged GluA1 (GluA1-SEP) and mRuby2. Cells were treated with DMSO as a control or CRT prior to the bleaching event. Images show dendritic spines before bleaching, as well as 5, 60 and 600s afterwards. (b) Fluorescence recovery curve and (c) one-phase association curve fitting of the GluA1-SEP intensity after bleaching. Data shown as mean ± SEM (b) or with 95% CI (c) from 3-4 independent experiments; n= 14 and 27 analyzed dendritic spines from 12 and 16 neurons, for DMSO and CRT-treated conditions, respectively. The significance of differences was analyzed by a multiple t-test, using the Holm-Sidak method. (d) Recovery half-time of the GluA1-SEP signal and (e) mobile GluA1 fraction. Data shown as mean ± 95% CI. (f) Representative inverted single-channel and colored merge pictures of neuronal dendritic branches expressing EGFP or caPKD-EGFP. Cells were treated with DMSO (-) or CRT, fixed and stained for the early endosome marker EEA1 and for GluA1. Arrows point to the EEA1-positive early endosomes included in the analysis. The EGFP signal is not included in the colored merged image. (g) Manders’ overlap coefficient for EEA1 and GluA1. Quantification of pictures shown in (f). The boxplot shows the results of three independent cultures. Center lines show the medians; box limits indicate the 25th and 75th percentiles; whiskers extend 1.5 times the interquartile range from the 25th and 75th percentiles, outliers are represented by dots; n = 28 cells each. The significance of differences was analyzed by a Kruskal-Wallis test with Dunn’s multiple correction. Representative Western Blot displaying total and pS845 GluA1 (h) as well as total and pS831 GluA1 (j) levels of DMSO (-) or CRT-treated hippocampal neuronal cultures. Actin served as a loading control. (i and k) Quantification of data shown in (h) and (j), respectively, using densitometry analysis. Data are presented as mean line density ± SEM of CRT normalized to the control (-), each dot represents one independent experiment (n=3). Statistical comparison by unpaired two-tailed t-test. The dotted line indicates the control level. *p<0.05, **p<0.01. Scale bar 2 μm

It is well established that constitutive endocytosis at perisynaptic zones regulates the number of AMPARs within the postsynaptic membrane (Ehlers 2000; Luscher *et al*. 1999; Rosendale *et al*. 2017; Man *et al*. 2000). Furthermore, under basal conditions, AMPARs undergo a robust time-dependent endocytosis (Ehlers 2000). To clarify whether PKD is involved in basal AMPAR endocytosis, we firstly investigated whether PKD short-term inhibition affects basal trafficking of AMPARs to early endosomes. To that end, neuronal cells expressing EGFP or caPKD-EGFP were treated with CRT or DMSO, fixed, permeabilized and stained for GluA1 and EEA1, a marker of early endosomes (Mu *et al*. 1995). EEA1 and GluA1 were present on dot-like structures distributed all over the dendritic branch (Figure 3f). This is in line with earlier reports showing that EEA1 participates in the endocytosis of AMPARs and is highly expressed in the postsynaptic compartment of hippocampal synapses (Selak *et al*. 2006; Selak *et al*. 2000). To assess whether the amount of GluA1 at early endosomes changes upon increased PKD activity, we compared the co-localization of GluA1 and EEA1 by analyzing the Mander’s coefficient (Figure 3g) in EGFP or caPKD-EGFP expressing neurons. Our data show that constitutively active PKD significantly enhances the amount of GluA1 at early endosomes. This was reverted to the control level when cells were treated with CRT for 10 minutes prior to fixation.

To corroborate our finding on AMPAR endocytosis, we studied the phosphorylation of GluA1 at S831 and S845 depending on PKD activity. S831 phosphorylation is mediated by CaMKII and Protein Kinase C (PKC), promoting the targeting of GluA1 to the PSD and increasing single channel conductance (Barria *et al*. 1997; Diering *et al*. 2016; Roche *et al*. 1996) whereas phosphorylation of S845 is mediated by Protein Kinase A (PKA), leading to an increased single-channel open probability and to targeting or retention of GluA1 at the cell surface (Banke *et al*. 2000; Man *et al*. 2007; Diering & Huganir 2018). Cells were treated with CRT or DMSO for 10 min, lysed and both the total and phosphorylated levels of GluA1 were measured via Western Blot (Figure 3h and j). Remarkably, while short-term inhibition of PKD did not modify basal pS845 GluA1 levels compared to the control (Figure 3i), our data show a significant increase of basal pS831GluA1 in CRT-treated cells when compared to the control (Figure 3k). These observations are in line with our previous results showing enhanced GluA1 levels at the PSD, slowed down GluA1 cell surface trafficking, and increased spontaneous network activity upon PKD inhibition. As S831-phosphorylated GluA1 subunits are enriched at the PSD (Diering *et al*. 2016), it further suggests that the occurrence of GluA1-containing AMPARs at the PSD is enhanced in PKD-inhibited cells. Our data thus show that PKD activity coordinates the localization and endocytic trafficking of GluA1-containing AMPARs under basal conditions.

### PKD regulates activity-mediated AMPAR endocytosis

Given that PKD activity promotes basal endocytotic AMPAR trafficking, we next investigated whether the kinase also plays a role in agonist-induced AMPAR endocytosis. To do so, we employed the biotinylation assay of receptor endocytosis using sulfo-NHS-SS-biotin (Ehlers 2000). Briefly, biotinylated hippocampal cultures were treated with CRT or DMSO, followed by 2 minutes of 100 μM AMPA-evoked stimulation of AMPARs while simultaneously blocking NMDA receptors with 50 μM D-AP5 (Figure 4a; treatment is designated as “AMPA”). Cells were incubated additionally for 10 min to allow for endocytosis before the remaining cell surface biotin was cleaved by a glutathione wash and cells were processed for analysis. In this way, GluA1 subunits remain biotinylated only within endosomes. In line with previous results, AMPA treatment enhanced GluA1 endocytosis as indicated by increased levels of internalized GluA1 in comparison to total GluA1, albeit not significantly (126%± 8% SEM compared to the control, Figure 4b, c). Strikingly, ligand-induced endocytosis of AMPARs was completely prevented when cells were pre-treated with CRT (72%± 13% SEM compared to the control). Notably, AMPA treatment also enhanced autophosphorylation of PKD, indicating that the kinase is a downstream target of AMPAR signalling. The AMPA-induced increase in kinase activity was completely blocked by pre-treatment of cells with CRT (Figure 4d, e).

**Fig. 4.**
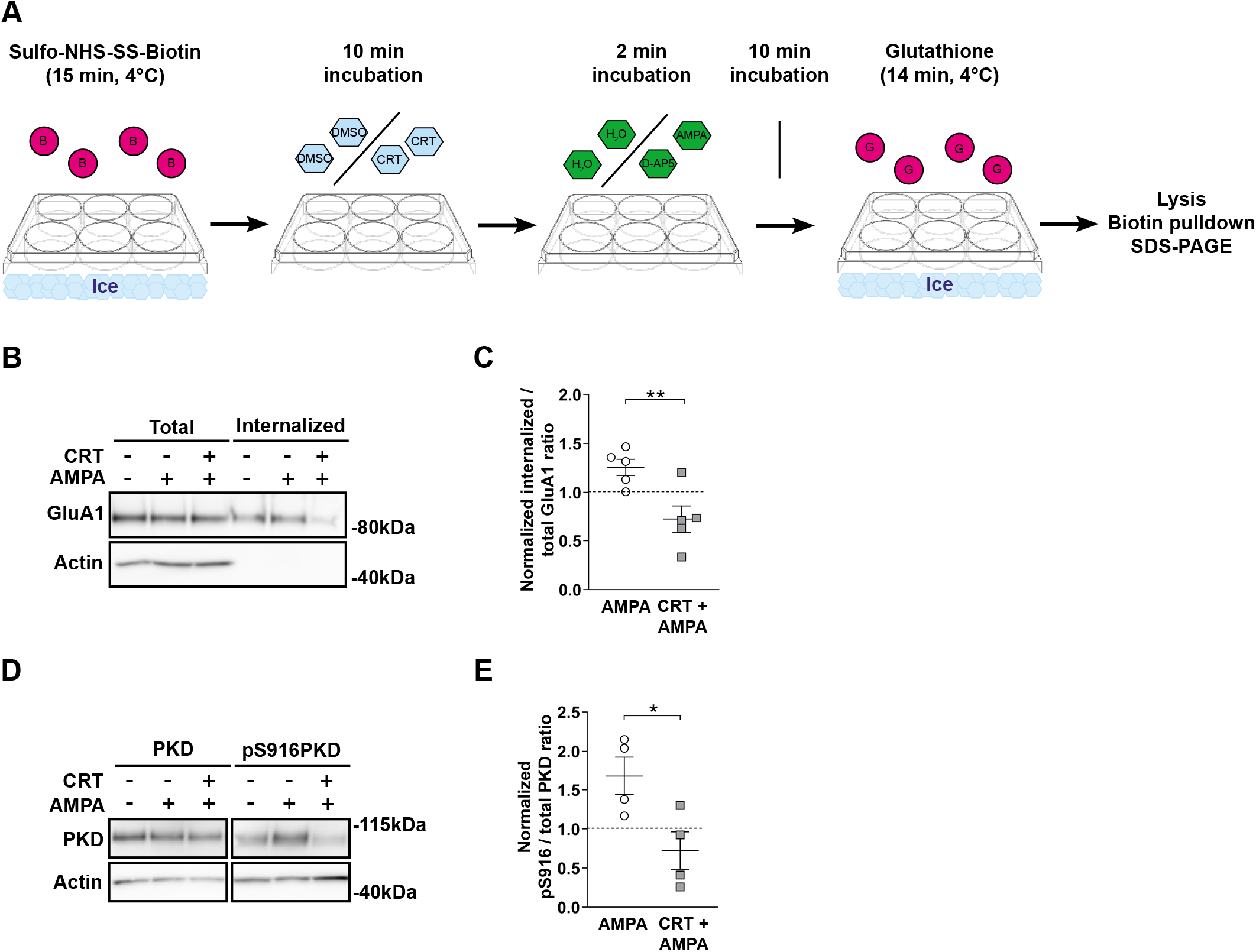
PKD activity is required for agonist-induced AMPAR endocytosis. (a) Scheme displaying the surface biotinylation assay for receptor internalization. After surface biotinylation, cells were treated with DMSO or CRT followed by AMPA and D-AP5 application. Afterwards, cultures were treated with glutathione to remove remaining surface biotin. Thereby, biotinylated GluA1 signal is detected only in endocytosed AMPARs, evoked by combined AMPA and D-AP5 treatment. Representative Western Blots displaying (b) total and internalized GluA1 levels and (d) PKD and pS916PKD levels in hippocampal neuronal cultures. Actin served as a loading control and is absent from the precipitated samples. (c, e) Quantification of data shown in (b, d), using densitometry analysis. Data are presented as mean line density ±SEM of the treated samples (AMPA and CRT plus AMPA) normalized to the control, each dot represents one independent culture. Statistical comparison was done by one-way ANOVA with Sidak’s correction. The dotted lines represent the control level.

Treatment of hippocampal neuronal cultures with NMDA promotes activity-dependent endocytosis of AMPARs thereby evoking LTD (designated as cLTD) (Beattie *et al*. 2000; Collingridge *et al*. 2010; Lee *et al*. 1998; Lee *et al*. 2002; Lin & Huganir 2007). In order to investigate whether PKD activity is also required for GluA1 internalization evoked by cLTD treatment, cells were pre-treated with CRT or DMSO for 10 min followed by cLTD treatment for additional 5 min. Cells were then changed to an NMDA-free medium and incubated for 15 min to allow for receptor endocytosis before undergoing surface protein biotinylation using sulfo-NHS-SS-biotin (Figure 5a). After cell lysis, biotinylated proteins were precipitated and the total expression and surface levels of GluA1 were detected through Western Blot analysis (Figure 5b). Remarkably, cLTD treatment promoted a significant decrease in surface GluA1 (78%± 1% SEM compared to the control) which could be prevented with CRT (124%± 10% SEM compared to the control, Figure 5c). Our data also show an enhanced PKD phosphorylation upon cLTD treatment indicating that PKD is activated downstream of NMDA receptors. In line with our previous data (Figure 1), CRT treatment completely prevented NMDA-induced PKD activation (Figure 5d, e). GluA1 dephosphorylation at S845 is a prerequisite for the LTD-induced endocytosis of AMPARs (Diering & Huganir 2018; Lee *et al*. 1998; Lee *et al*. 2003). In line with the literature, cLTD treatment promoted a significant decrease in the level of pS845 GluA1 compared to the control. Notably, the NMDA-induced dephosphorylation of S845 was not blocked with CRT treatment (Figure 5f, g) suggesting that PKD acts further downstream.

**Fig. 5.**
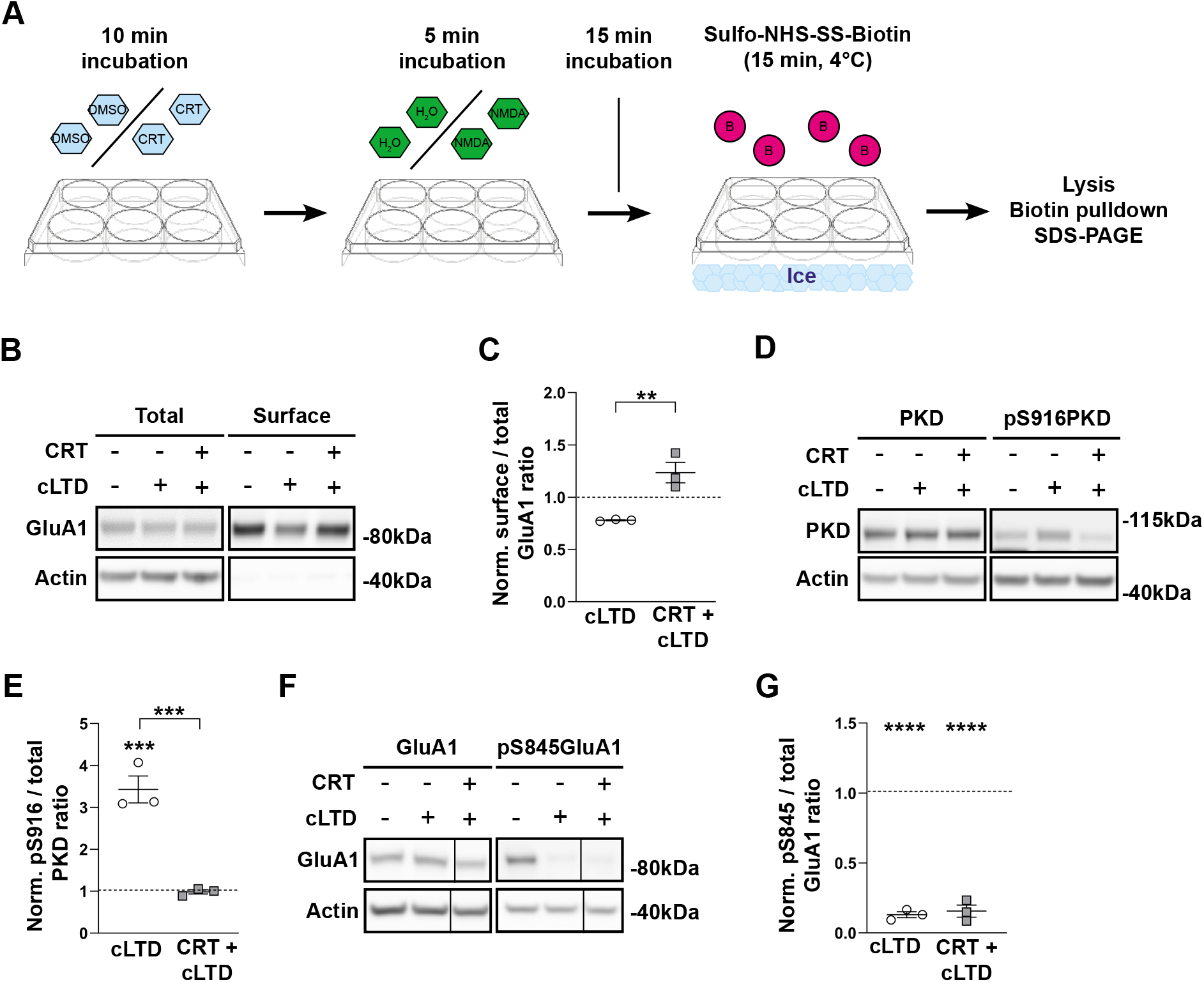
PKD activity is required for cLTD-induced AMPAR endocytosis. (a) Scheme displaying the cell surface biotinylation assay to detect the remaining surface GluA1 receptors after cLTD treatment. Cells were treated with DMSO or CRT followed by NMDA application. After the cLTD treatment, cultures were surface biotinylated using sulfo-NHS-SS-biotin and further processed. Representative Western Blots displaying (b) total and surface GluA1 levels, (d) PKD and pS916PKD levels, and (f) total and pS845GluA1 levels in biotinylated hippocampal neuronal cultures. Actin served as a loading control and is absent from the precipitated samples. (c, e, g) Quantification of data shown in (b,d,f), using densitometry analysis. Data are presented as mean line density ± SEM of treated samples (cLTD and CRT plus cLTD) normalized to the control; each dot represents one independent culture. Statistical comparison was done by one-way ANOVA with Sidak’s correction. The dotted lines represent the control level. The boxplots show the results of three independent cultures. Center lines show the medians; box limits indicate the 25th and 75th percentiles; whiskers extend 1.5 times the interquartile range from the 25th and 75th percentiles, outliers are represented by dots. The significance of differences was analyzed by a Mann-Whitney test (two-tailed). **p<0.01, ***p<0.001, ****p<0.0001.

MEA recordings were additionally taken and compared between control and cLTD treated cultures. Our prior experiments in patch clamp settings have already shown that firing output of such cultures are governed by potent depolarizing events when large numbers of neuron fire synchronously and drive the activity of postsynaptic cells via glutamatergic connections (network bursting)(Ratkai *et al*. 2021). In line with the reduced AMPA receptor levels upon cLTD treatment, network activity was significantly decreased, as indicated by elevated interspike interval medians (Figure 6a, b). In addition, bursts were completely diminished upon cLTD as also indicated by the extremely elevated ISI medians normalized to pre-cLTD values (Figure 6c). This latter observation indicates the reduction of potent depolarizing events, compound excitatory postsynaptic potentials that are the main contributors of action potential emissions of cultured neurons. Importantly, when CRT was present during NMDA treatment and for an additional 10 min, normalized interspike intervals were increased to a significantly lower extent compared to DMSO treated cultures (Figure 6c).

**Fig. 6.**
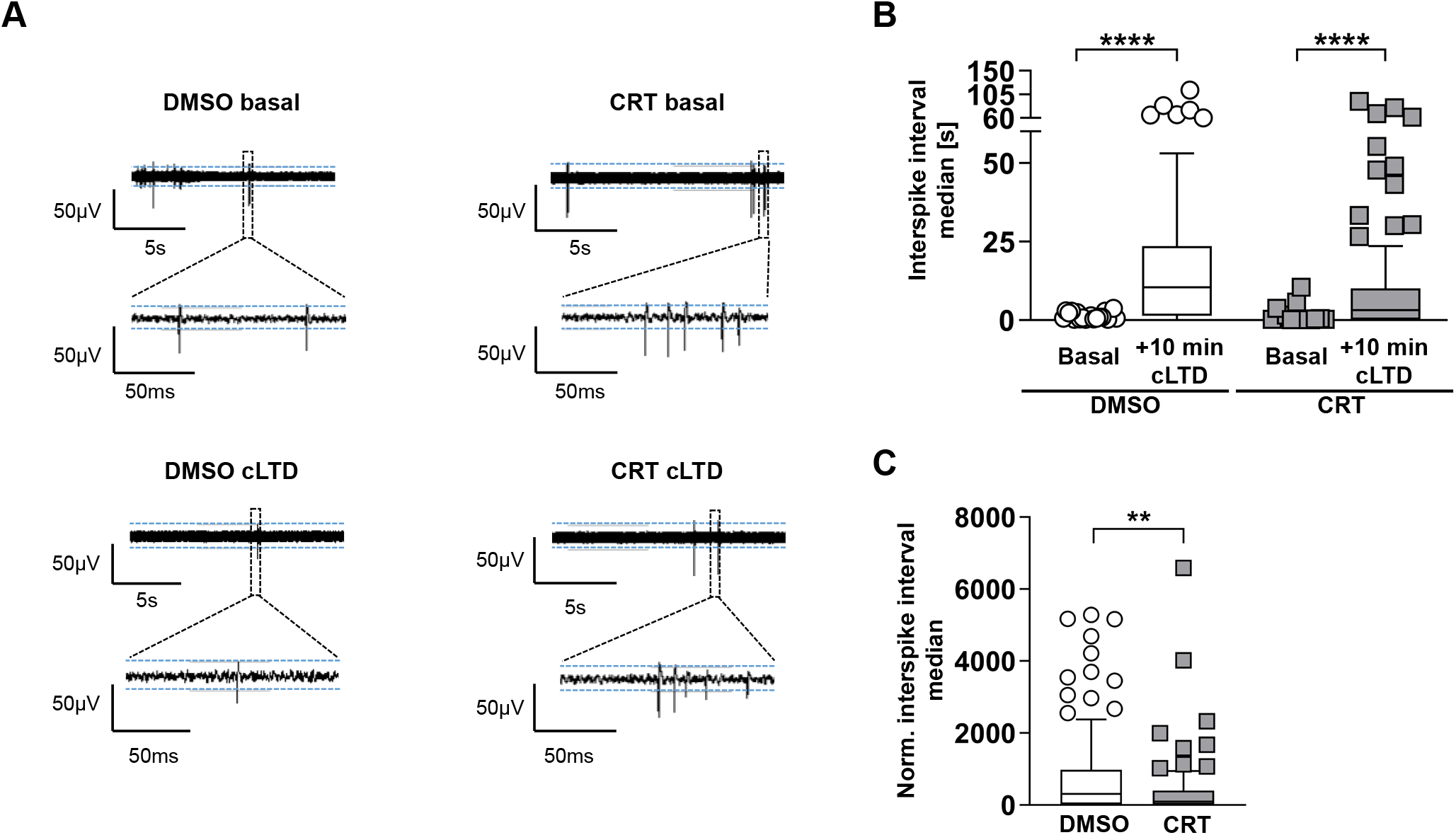
PKD activity controls neuronal network activity. (a) Short segments of raw voltage traces before and after CRT treatment und basal conditions (top) and upon cLTD (bottom). (b) Quantification of the raw interspike interval median of neuronal cultures both before and after treatment with DMSO or CRT for 10 minutes, followed by treatment with NMDA and a subsequent 10 min incubation. (c) Interspike interval median of cells treated with 10 min of DMSO or CRT followed by NMDA application and a subsequent incubation period of 10 min. Data were normalized for their pre-treatment values. The boxplots show the results of three independent cultures. Center lines show the medians; box limits indicate the 25th and 75th percentiles; whiskers extend 1.5 times the interquartile range from the 25th and 75th percentiles, outliers are represented by dots. The significance of differences was by a Wilcoxon matched-pairs signed rank test (two-tailed) (b) or by a Mann-Whitney test (two-tailed) (c). *p<0.05, **p<0.01, ***p<0.001, ****p<0.0001.

Collectively, our data demonstrate that PKD activity contributes to agonist- and activity-induced endocytosis of AMPARs and is required for a physiological change in network activity.

## Discussion

In neurons, PKD regulates Golgi function and polarized secretory trafficking thereby contributing to dendrite development and the maintenance of neuronal polarity (Czondor *et al*. 2009; Quassollo *et al*.2015; Horton *et al*. 2005). Moreover, PKD activity is required for memory formation and learning (Bencsik *et al*. 2015). However, a role for PKD in the turnover of AMPARs has not been described yet. In the present study, we show that short-term inhibition of PKD activity increases the level of surface and synaptic GluA1-containing AMPARs under basal conditions and during chemically evoked AMPAR internalization. We further demonstrate that PKD is activated downstream of NMDA and AMPA receptor signaling and promotes AMPAR endocytosis.

The dynamic regulation of the number of synaptic AMPARs is one of the main mechanisms modifying long-term synaptic efficacy. Our data highlight PKD as a novel member of the postsynaptic signaling machinery regulating basal and activity-induced endocytosis of AMPARs. This is in line with our previous findings that PKD activity is required for the maintenance of the dendritic tree (Czondor *et al*. 2009), as well as for the stabilization of the F-actin network within dendritic spines upon induction of LTP and learning and memory formation *in vivo* (Bencsik *et al*. 2015). Accordingly, a dual role for PKD in synaptic plasticity becomes apparent: on the one hand, the kinase controls the activity-induced potentiation and enlargement of spines during chemically-induced LTP through stabilization of the F-actin network within the newly enlarged dendritic spines (Bencsik *et al*. 2015), while on the other hand it is required for the elimination of AMPARs from the cell surface during cLTD.

In line with our results, PKD has been already described to participate in endocytosis and recycling of membrane proteins in non-neuronal cells. In fibroblasts, PKD activity is necessary for PDGF-driven short-loop recycling of α_v_β_3_ integrin (Woods *et al*. 2004). However, due to its established role in polarized secretory trafficking in neurons (Horton *et al*. 2005), inhibition of PKD activity is expected to affect exocytotic trafficking as well. Our FRAP data show that upon short-term PKD inhibition, GluA1-SEP signal recovered significantly slower, albeit to the same level as the control. This suggests that the mobile fraction of GluA1-SEP was unaltered, while PKD inhibition slows the trafficking of cell surface GluA1-SEP. In the absence of activity, recombinant GluA1-SEP is largely mobile on spines and the fast GluA1-SEP recovery mostly depends on lateral diffusion of plasma-membrane resident AMPARs from non-synaptic sites while exocytosis only has a small contribution (Makino & Malinow 2009). Therefore, based on our observation that caPKD expression significantly decreased the amount of GluA1 at the cell surface while increasing the amount of GluA1 in early endosomes and that the PKD inhibitor CRT prevented this, we hypothesize that PKD inhibition impairs the internalization of bleached GluA1-SEP, thereby slowing the rate at which non-synaptic, unbleached GluA1-SEP enters the postsynaptic membrane (Shi *et al*. 1999; Opazo & Choquet 2011; Opazo *et al*.2012; Lisman & Raghavachari 2006). However, we cannot fully exclude that short-term PKD inhibition might affect lateral mobility of AMPARs as well.

The small GTPase Rab5 regulates the fusion kinetics of plasma-derived endocytic vesicles with early endosomes in both basal and activity-mediated trafficking of AMPARs (Bucci *et al*. 1992; Gorvel *et al*. 1991; Sziber *et al*. 2017) and is essential for LTD (Brown *et al*. 2005). In accordance, loss or inhibition of the Rab5 effector protein EEA1 results in increased GluA1-containing AMPAR surface expression (Selak *et al*.2006; Xu & Pozzo-Miller 2017) reminiscent of the increased GluA1 surface expression observed upon short-term PKD inhibition. Additionally, expression of constitutive active PKD enhanced the co-localization of GluA1 with EEA1, supporting a role for PKD in promoting AMPAR endocytosis through regulating Rab5 activity. To date, two PKD substrates, RIN1 and Rabaptin-5, are known to be involved in the endocytosis of surface receptors. RIN1 is a Rab5 guanosine exchange factor (GEF) and we recently showed that its ability to enhance Rab5 activity is a critical step during activity-dependent AMPAR internalization (Sziber *et al*. 2017). PKD-mediated phosphorylation of RIN1 modulates its ability to activate Abl kinases (Ziegler *et al*. 2011), however, an effect on Rab5 activation and AMPAR endocytosis has not been investigated yet. Rabaptin-5 forms a complex with Rabex-5, a Rab5-GEF, and Rab5, thereby promoting Rab5 activation during endocytosis (Horiuchi *et al*. 1997; Zhang *et al*. 2014). Notably, PKD-mediated Rabaptin-5 phosphorylation is implicated in the regulation of integrin receptor trafficking in cancer cells (Christoforides *et al*. 2012). It is therefore intriguing to speculate that signaling from PKD to RIN1 and/or Rabaptin-5 may promote activation of Rab5 and thus AMPAR endocytosis.

The presence and regulation of post-translational modifications on neurotransmitter receptors modifies their function and trafficking and thus synaptic strength. Post-translational modifications have been described to mediate numerous processes including AMPAR membrane targeting, retention, conductance, endocytosis and degradation (Boehm *et al*. 2006; Chung *et al*. 2000; Coultrap *et al*. 2014; Esteban *et al*. 2003; Hayashi & Huganir 2004; Lee *et al*. 2010; Roche *et al*. 1996; Seidenman *et al*.2003; Steinberg *et al*. 2006; Widagdo *et al*. 2015). Several phosphorylation sites on GluA1 have already been shown to be important for stability at the PSD and AMPAR conductance (Boehm *et al*.2006; Derkach *et al*. 1999; Esteban 2003; Roche *et al*.1996), with S831 and S845 being best understood. CaMKII/PKC-mediated phosphorylation of S831 increases single-channel conductance (Barria *et al*. 1997; Derkach *et al*. 1999). Moreover, under basal conditions, pS831-GluA1 receptors are enriched at the PSD suggesting that phosphorylation at S831 increases GluA1 targeting to this compartment (Diering *et al*. 2016). In agreement with this, we detected an increase in pS831 GluA1 levels upon short-term PKD inhibition, presumably as a consequence of impaired AMPAR endocytosis. PKA-mediated phosphorylation of S845 has been described to promote GluA1 targeting to the cell surface and single-channel open probability (Banke *et al*. 2000; Lee *et al*. 2010; Man *et al*.2007). Notably, dephosphorylation at S845 is required for AMPAR endocytosis upon LTD (Lee *et al*.2010). Our data show that acute CRT treatment did not prevent the dephosphorylation of GluA1 at serine 845 in response to cLTD. This suggests that i) PKD activity promotes AMPAR endocytosis downstream of S845 dephosphorylation and ii) upon PKD inhibition, AMPARs are primed for their endocytosis but remain on the surface.

Based on our data we propose the following model on how PKD regulates AMPAR endocytic trafficking: under basal conditions, endogenous PKD is active and promotes the removal of AMPARs from the synaptic membrane by Rab5-dependent endocytosis. Chemically-induced AMPAR endocytosis either through NMDA receptor or AMPAR stimulation further enhances PKD activity thus accelerating AMPAR elimination from the surface.

As most excitatory transmission in the brain is mediated by the AMPARs, their dysfunction has been observed in many neuronal disorders, with Alzheimer disease (AD) being the best reported so far. High concentration of soluble oligomeric Aβ induces the removal of surface AMPARs at synapses, leading to synaptic depression and impaired synaptic plasticity and memory (Hsieh *et al*. 2006; Shankar *et al*.2007). Strikingly, at preclinical stages of AD, the appearance of enlarged Rab5-positive early endosomes was associated with Rabaptin-5 translocation to endosomes (Cataldo *et al*. 2000). Indeed, hyperactivation of Rab5 has been observed in AD patients and mouse models of AD disease (Ginsberg *et al*. 2010a; Ginsberg *et al*. 2011; Ginsberg *et al*. 2010b). These data imply that endocytic uptake of AMPARs is enhanced in AD, however, whether this also correlates with increased PKD activity needs to be addressed in future studies.

## Acknowledgements

We are grateful to Viktória Szentgyörgyi and Gisela Link (Institute for Cell Biology and Immunology, Germany) for their excellent technical assistance and Raluca Tamas (Institute for Cell Biology and Immunology, Germany) for critical reading of the manuscript. The manuscript has been deposited as a preprint in bioRxiv (doi: https://doi.org/10.1101/2020.04.09.033514).

## Authors’ contributions

**Conceptualization:**Angelika Hausser, Katalin Schlett; **Methodology:**Carlos O. Oueslati Morales, Attila Ignacz, Norbert Bencsik, Aniko E. Ratkai, Wolfgang S. Lieb; **Formal analysis and investigation:**Carlos O. Oueslati Morales, Attila Ignacz, Attila Szücs, Aniko E. Ratkai, Norbert Bencsik, Stephan A. Eisler, Wolfgang S. Lieb; **Writing - original draft preparation:**Carlos O. Oueslati Morales; **Writing - review and editing:**Angelika Hausser, Katalin Schlett; **Funding acquisition:**Angelika Hausser, Katalin Schlett. All authors contributed to the article and approved the submitted version.

## List of Abbreviations

AD: Alzheimer’s Disease
AMPA: α-amino-3-hydroxy-5-methyl-4-isoxazolepropionic acid
AMPAR: AMPA receptor
BCP: Burst Cycle Period
CaMKs: Calcium/calmodulin-dependent protein kinases
caPKD: constitutively active PKD
CI: Confidence Interval
cLTD: chemical LTD
CRT: CRT0066101
DIV: Days In Vitro
EEA1: Early Endosome Antigen 1
EGFP: Enhanced Green Fluorescent Protein
FBS: Fetal Bovine Serum
FRAP: Fluorescence Recovery After Photobleaching
GEF: Guanosine Exchange Factor
ISI: Inter Spike Time Interval
kdPKD: kinase dead PKD
LTD: Long-term depression
LTP: Long-term potentiation
MEA: Multi-Electrode Array
PKA: Protein kinase A
PKC: Protein kinase C
PKD: Protein kinase D
pS831GluA1: phospho-serine 831 GluA1
pS845GluA1: phospho-serine 845 GluA1
pS916PKD: phospho-serine 916 PKD
PSD: Post Synaptic Density
GluA1-SEP: Super Ecliptic pHluorin GluA1

## Footnotes

1 ResearchGate. (2018). NeuroExpress: Analysis software for whole-cell electrophysiological data. https://www.researchgate.net/proiect/NeuroExpress-Analysis-software-for-whole-cell-electrophysiological-data. [Accessed October 26, 2020].

